# Identification of a Thermogenic Target in the Dorsal Raphe Nucleus for Weight Management

**DOI:** 10.1101/2025.04.24.650283

**Authors:** Alexandre Moura-Assis, Kaja Plucińska, Aiste Baleisyte, David Meseguer, Bandy Chen, Kyle Pellegrino, Luca Parolari, Jordan T. Shaked, Keith J. Page, Douglas W. Barrows, Thomas S. Carroll, Jose G. Grajales-Reyes, Daxiang Na, Hans H. Schiffer, Yun Chen, Huikai Sun, Nidhi Kaushal, Holger Monenschein, Daniel F. Barker, María José Ortuño, Nicolas Renier, Paul Cohen, Mark Carlton, Nathaniel Heintz, Nicola L. Brice, Jeffrey M. Friedman, Marc Schneeberger

## Abstract

Obesity emerges from a complex interplay of factors, including imbalanced interoception, genetic predisposition, and environmental cues, ultimately disrupting body weight homeostasis^1^. While much research has concentrated on strategies to suppress appetite for sustained weight loss, insufficient attention has been given to counterregulatory mechanisms that promote energy expenditure. Here, we show that chronic inhibition of GABAergic neurons in the Dorsal Raphe Nucleus (DRN^VGAT^) reduces body weight in diet-induced obese (DIO) mice. In this study, molecular profiling and *in-situ* hybridization in rodent and human brains revealed that the constitutively activated orphan receptor GPR6 is selectively enriched in DRN^VGAT^ neurons. We next developed and administered a potent and highly selective GPR6 inverse agonist, which significantly reduced weight gain in DIO mice by stimulating brown adipose tissue thermogenesis without affecting appetite. Altogether, this study transitions from transcriptomic profiling, high-throughput drug screening and metabolic phenotyping to successfully identify a novel candidate to treat obesity.

## Main

Overweight and obesity are highly prevalent and currently affect a staggering 38% of the global population^2^. Particularly alarming is the surge in childhood obesity, which may have lasting effects on neurodevelopment^3,4^. The intricate brain circuits orchestrating feeding behaviors and energy expenditure are particularly affected in this context^5^. Readily available calorie-dense food exacerbates the imbalance between energy supply and demand, imprinting neural circuits and resetting homeostatic baselines^6,7^. While recent pharmacological interventions to suppress appetite have been clinically promising, they often have undesirable side effects and tend to plateau in efficacy^8–13^.

Consumption of a high-fat diet (HFD) rewires homeostatic and hedonic neural systems involved in feeding and body weight regulation. For example, agouti-related peptide (AgRP)-expressing neurons in the hypothalamus are pivotal in regulating feeding behavior and are profoundly affected by HFD intake^14^. DIO blunts the inhibition of AgRP neurons by exteroceptive cues, such as food presentation, and interoceptive signals, such as intragastric fat infusion^15^.

Building on these studies, we hypothesized that a relatively short exposure to HFD could alter the activity patterns of distinct brain regions before the onset of obesity in mice. We employed an unbiased screen based on the whole-brain expression of the immediate early gene Fos, using iDISCO+ and ClearMap^16^. Among the regions significantly more active in mice fed a HFD for 7 days compared to chow-fed mice, two clusters of regions drew our attention: (1) hypothalamic nuclei, such as the paraventricular hypothalamus and lateral hypothalamus and (2) brainstem nuclei, predominantly the DRN **(Figure 1A and Extended Data Fig. 1A)**. Given our prior work in the DRN with regards to energy balance regulation^17,18,19^, we next tested whether DRN^VGAT^ neurons are persistently activated in DIO mice which, if true, would support our hypothesis that inhibiting them would reduce body weight in obese mice. Toward this end, we quantitated the expression of cFOS in the DRN of mice fed a HFD for 3 days (acute exposure), 2 weeks (prolonged exposure), and 16 weeks (chronic exposure) and found a significant increase in cFOS-positive neurons in the DRN at all time points, indicating heightened activity induced by HFD consumption **(Figure 1B)**. We next performed fluorescent *in-situ* hybridization (RNAscope) studies to quantify the percentage of cFOS-positive neurons that aligned with *slc32a1* (vGat) in the DRN and found co-expression of vGat in approximately 25% of the activated cells **(Extended Data Fig. 1B).**

**Figure 1.**
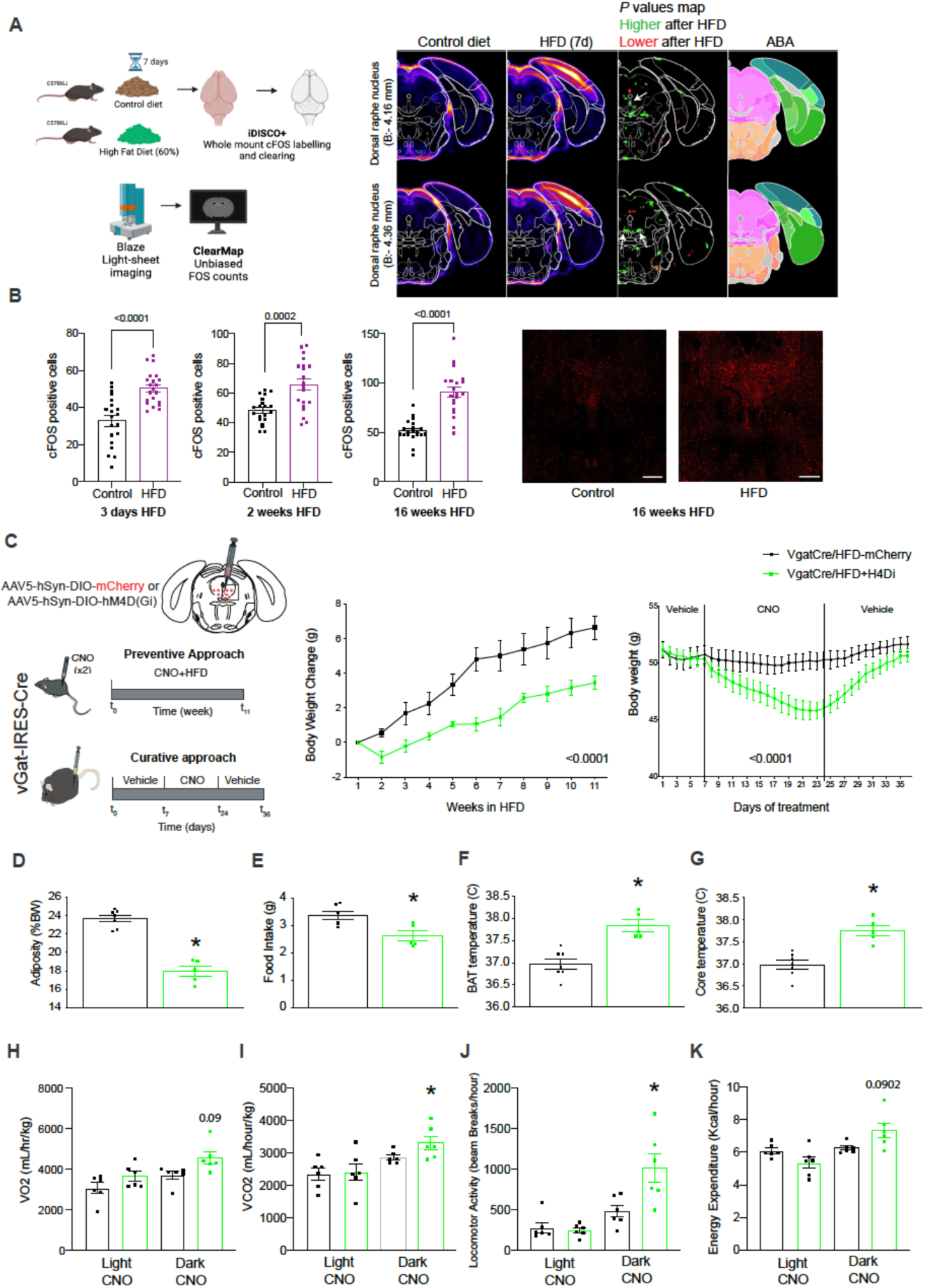
Chronic inhibition of DRN^VGAT^ neurons ameliorates obesity. (A) Schemata (created with BioRender.com) to map activity-dependent Fos expression in mice fed a HFD for 7 days. Right panel: coronal views of the p-value of the voxel maps for Fos-positive cell densities at two different levels of DRN. White arrows indicate DRN regions significantly activated after 7 days of HFD feeding. ABA (Allen Brain Atlas) annotated image. (B) Time-course quantification of cFOS-positive neurons in the DRN of mice fed an HFD for 3 days, 2 weeks, and 16 weeks and representative immunohistochemical staining of cFOS in the DRN of mice fed an HFD for 16 weeks. (scale bar = 100um) (C) Schemata for chemogenetic inhibition of DRN^VGAT^ neurons in vGat-IRES-Cre mice injected with CNO twice a day for 11 weeks in the preventive approach or injected daily for 17 days in DIO mice for the curative approach (right). Body weight change in mice fed a HFD for 11 weeks along with DRN^VGAT^ neurons chemogenetic inhibition (n= 6-7 per group, *P* <0.0001) (middle) and body weight of DIO mice before and after chemogenetic inhibition of DRN^VGAT^ neurons for 17 days (right) (n= 6-7 per group, *P* <0.0001). (D-G) (n= 6-7 per group, P < 0.05), Metabolic phenotyping of vGat-IRES-Cre after completion of the preventive approach. (D) Adiposity (% total body weight) (E) Average food intake (F) Interscapular BAT temperature (G) Whole-body median temperature (H-K) Indirect calorimetry of DIO Vgat-IRES-Cre mice injected inhibitory DREADDs and placed in metabolic cages for measurement of (H) oxygen consumption (VO_2_), oxygen dioxide production (VCO_2_), (J) locomotion activity and (K) energy expenditure. Data are represented as mean ± s.e.m. *P* values were calculated using a or an unpaired two-tailed Student’s *t*-test (**A, D-G**), two-way ANOVA with a multiple-comparisons test using a Tukey post-hoc approach (**C**) or using CalR software and a two-sided ANCOVA regression analysis taking body weight into account (**H-K**). *P* < 0.05 is considered significant. The CNO dose used was 1 mg/kg.

Since activating these neurons increases body weight, we reasoned that blunting the overactivation of DRN^VGAT^ neurons would prevent mice from gaining excess body weight when fed a HFD. To test this, we first turned to chemogenetics and injected an AAV5-hSyn-DIO-hM4D(Gi) virus into the DRN of vGat-Cre mice and treated experimental and control mice (AAV5-hSyn-DIO-mCherry) with clozapine-N-oxide (CNO) at the same time as the mice were switched from chow to HFD. We found that chemogenetic inhibition of DRN^VGAT^ neurons effectively suppressed body weight gain compared to control littermates **(Figure 1C)**. Similarly, treatment with CNO also significantly reduced body weight and adiposity in obese mice fed a HFD for 16 weeks **(Figure 1C – right panel and D)**. This inhibition also led to a significant decrease in food intake **(Figure 1E)** and an increase in energy expenditure, evidenced by elevated BAT thermogenesis **(Figure 1F)**, core body temperature **(Figure 1G)**, indirect calorimetry assessment **(Figure 1H, I and K)** and locomotor activity **(Figure 1J)**.

Notably, after the removal of CNO treatment, we observed that mice started to regain body weight when injected with vehicle, suggesting that chronic inhibition of DRN^VGAT^ neurons is necessary to maintain weight loss (**Figure 1C** – right panel). A recent study has shown that DRN^VGAT^ neurons are activated during feeding-related behaviors, such as food contact, and their responsiveness to food cues is increased by hunger and palatability^20^. Behavioral screening experiments also demonstrated that DRN^VGAT^ neurons’ activation increased contact duration with an inedible object, reduced walking duration, and decreased chamber exploration, suggesting a possible role for DRN^VGAT^ neurons in maintaining food consumption^20^.

The heightened activation of DRN^VGAT^ neurons observed during the consumption of highly palatable food and the significant real-time place preference for the light-paired chamber elicited by optogenetic activation of these neurons^21^ raises intriguing possibilities about their potential involvement in rewarding-promoting behaviors. However, the specific reward circuitry by which DRN^VGAT^ neurons operate remains to be determined.

Our results showing that chemogenetic inhibition of DRN^VGAT^ neurons reduces the body weight of DIO mice suggest that small molecules inhibiting their activity would also reduce weight. We thus set out to identify druggable targets enriched in DRN^VGAT^ neurons that could be pharmacologically targeted to inhibit these neurons. We profiled these neurons by introducing a virus expressing a Cre-dependent GFPL10a fusion protein into the DRN of vGat-Cre mice. We then used translating ribosomal affinity purification (TRAP) to molecularly profile DRN^VGAT^ neurons **(Figure 2A),** followed by differential expression analysis of genes enriched in the immunoprecipitated (IP) RNA over RNA input **(Figure 2B)**. These data identified 24,100 genes, of which 2,257 genes were significantly enriched (1.5-fold-enrichment with an adjusted P value (p<0.05) **(Supplementary Table 1)**. The molecular phenotyping of DRN^VGAT^ neurons was subsequently compared with neuronal transcriptomes available in the GENSAT database^22^. 8,519 genes were enriched in DRN^VGAT^ compared with other neural populations cataloged in the GENSAT database **(Supplementary Table 2)**. To identify druggable targets, we used the Gene Ontology browser^23^ database to identify genes enriched in DRN^VGAT^ that were also listed in both the signaling receptor activity <u>(GO:0038023)</u> and plasma membrane <u>(GO:0005886)</u> categories (<u>MGI,</u> informatics.jax.org) **(Figure 2C)**. After applying these filters, 259 druggable receptors were identified with more than 1.5-fold enrichment in DRN^VGAT^ neurons compared to the GENSAT database (highlighted in **Supplementary Table 3; Supplementary** Figure 2). Next, we evaluated the expression patterns of the enriched genes using the Allen Brain Atlas database^24^ to identify the most differentially expressed receptors in the DRN^VGAT^ neurons compared with other cell types. Three orphan G-protein-coupled receptors (*gpr3, gpr6,* and *gpr39*) showed significantly higher expression in the DRN **(Figure 2C)**. Consistent with this, aggregate differential expression analysis and the relative expression of these selected receptors, compared with molecular profiles of other cell types in the GENSAT database, reconfirmed the cell-specific enrichment of these genes in DRN^VGAT^ neurons **(Figure 2D)**. (TRAP IP vs input: *gpr3*: 2.2-fold enrichment, *gpr6*: 5.7-fold enrichment and *gpr39*: 2.5-fold enrichment; relative expression to GENSAT: *gpr3*: 2.6-fold enrichment, *gpr6*: 21-fold enrichment and *gpr39*: 200-fold enrichment). Of further interest, sphingosine 1-phosphate is a potential ligand for GPR3 and GPR6 and has a cannabinoid-like structure^25^. Moreover, its plasma levels are elevated in obesity^26^. GPR39, in turn, is a Zinc signaling receptor^27^. We decided to focus on GPR6 because is almost exclusively expressed in the central nervous system and is more highly enriched in DRN than GPR3 and GPR39.

**Figure 2.**
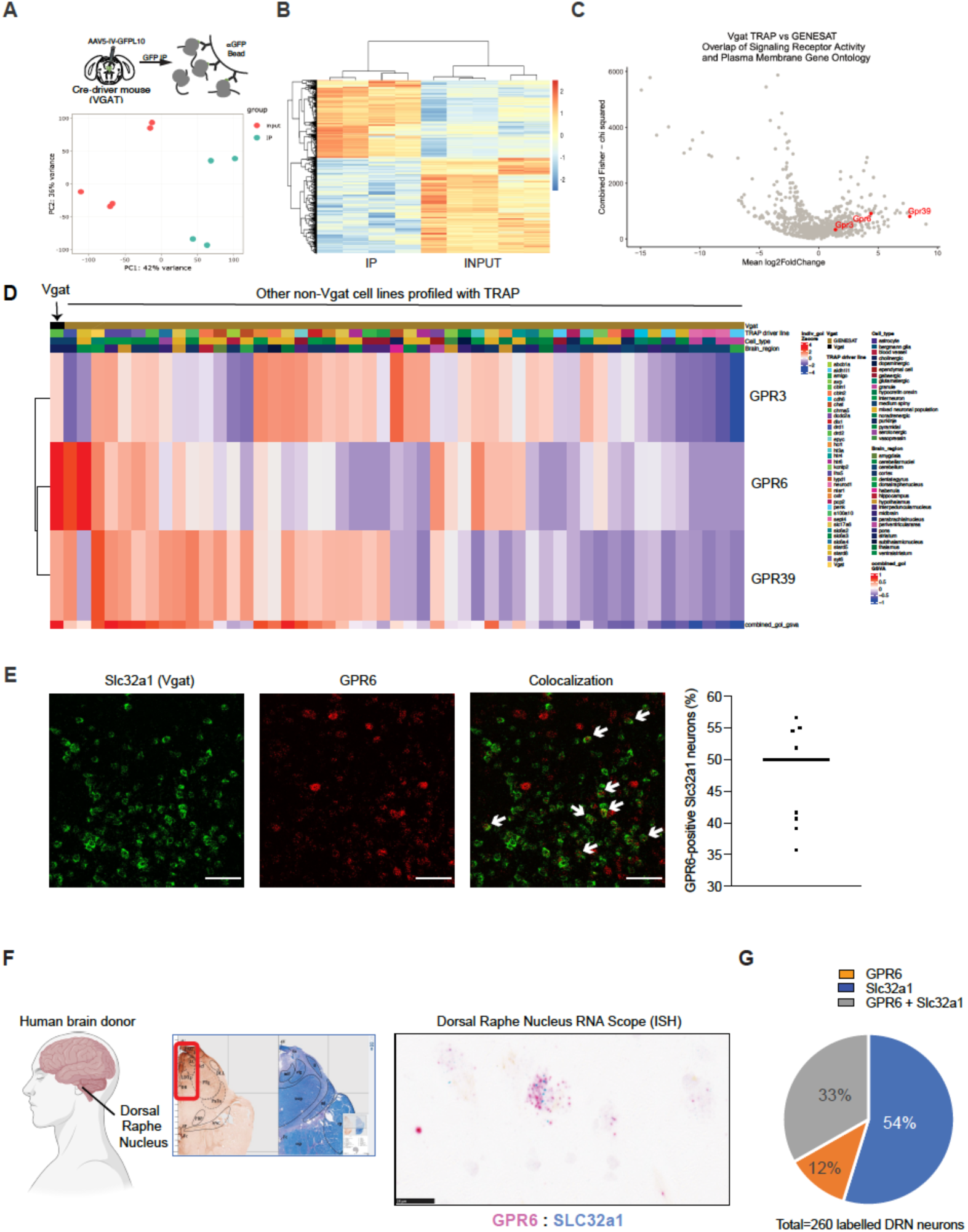
Genetic profiling of DRN^VGAT^ neurons and GENSAT database comparison reveals GPR6 as a druggable target. (A) Schemata for viral TRAP after injection of AAV5-IV-GFPL10a in the DR of vGat-IRES-Cre mice. GFPL10a fusion protein is translated after Cre-mediated recombination and the ribosomal protein L10a tagged with GFP is immunoprecipitated (IP). PCA analysis of input and IP samples (bottom). (B) Heat map representing the comparison between 4 IP samples and 5 INPUT RNA samples (C) Volcano plot depicting the comparison between IP samples from our viral TRAP with RNA sequencing samples found in the GENSAT database. Plasma membrane and overlap of signaling receptor activity Gene Ontology filters were applied for identifying putative targets (highlighted in red). (D) Heatmap representing the expression levels of GPR3, GPR6 and GPR39 obtained from viral TRAP of DRN^VGAT^ neurons in comparison to immunoprecipitated samples from TRAP experiments available in the GENSAT database. (E) Fluorescent *in-situ* hybridization (RNAscope) photomicrographs of Slc32a1 (green) and GPR6 (red) in the DRN of wild-type mice displaying an average of 50% colocalization (n=9 samples, scale bar = 100um) (F) Representative *in-situ* hybridization photomicrographs of SLC32A1 (blue) and GPR6 (pink) in the DRN of human brain (scale bar= 25um) (G) Out of 260 neurons identified in human DRN, approximately 33% displayed overlap between Slc32a1 and GPR6.

We next confirmed that DRN neurons expressing GPR6 also express vGat using fluorescent *in-situ* hybridization (RNAscope) and found colocalization of GPR6 with vGat (*slc32a1*) in approximately 50% of *slc32a1*-positive neurons **(Figure 2E)**. Since our goal was to develop druggable strategies for weight loss in humans, we also obtained brain sections from human brain donors and found GPR6 expression in the human DRN **(Figure 2F)**. Quantitative assessment was performed using the Paxinos Atlas of the human brainstem following the ventral extension of the midline, revealing a 33% overlap between SLC32A1 and GPR6 **(Figure 2G)**. Together, these results demonstrate co-expression of these genes in human DRN^VGAT^ neurons, recapitulating our *in-situ* experiments in mice. GPR6 is a Gs-coupled receptor, suggesting that blocking this receptor could inhibit DRN^VGAT^ neurons and potentially provide a novel therapeutic approach for treating obesity in obese individuals.

While some data have suggested that S1P might be a ligand for GPR6, this has not been confirmed. As an alternative, an inverse agonist screen was used to identify CVN527, a novel, potent, and selective GPR6 inverse agonist developed by structure-activity analysis after a lead compound was identified in a high-throughput screen^28^ (**Figure 3A**).

**Figure 3.**
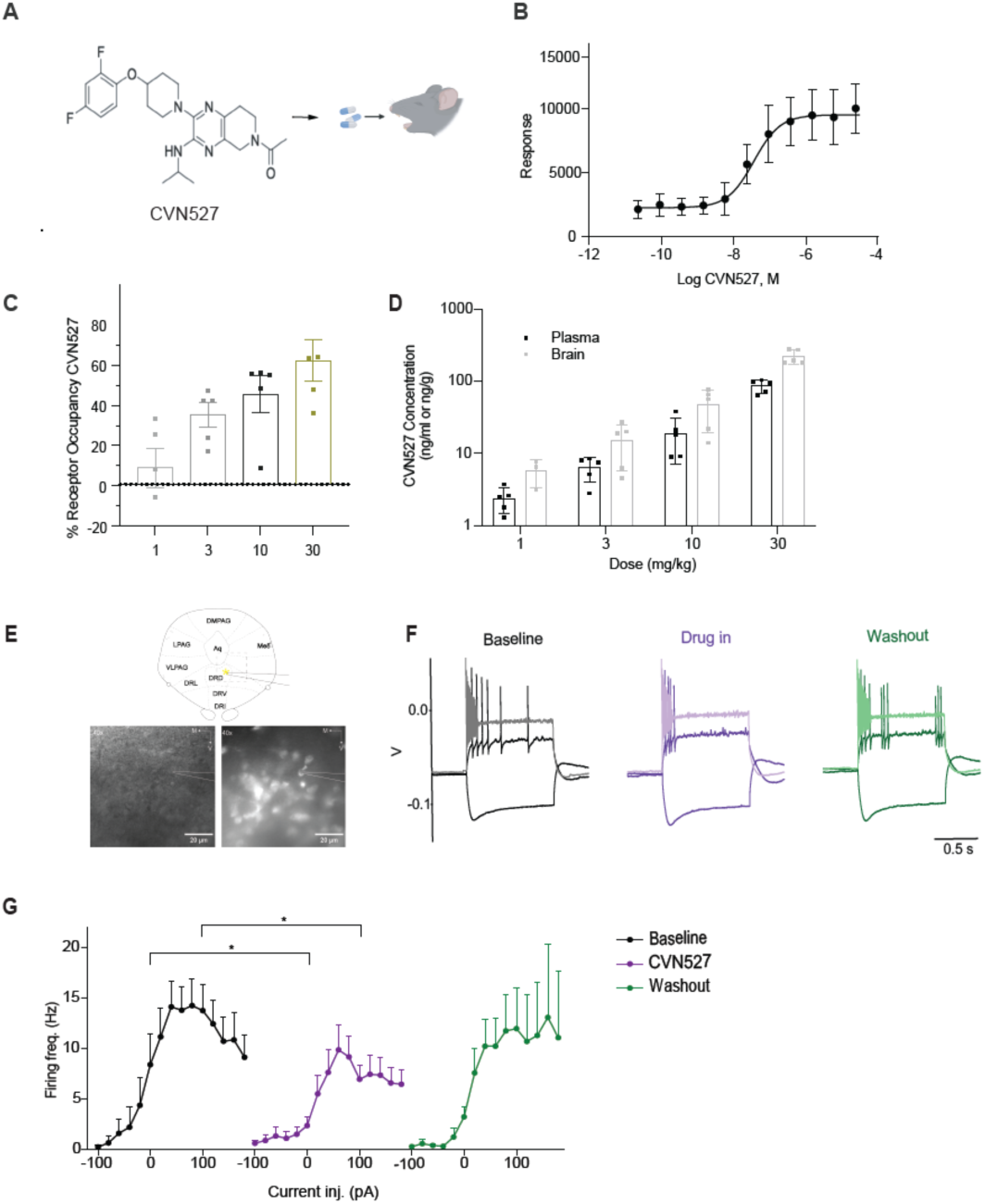
Development and electrophysiological validation of CVN527 as a potent, selective GPR6 inverse agonist. (A) Chemical structure of CVN527. (B) CHO-K1 cell transfected with mouse GPR6 cDNA. Results represent means of 3 independent experiments +/-S.D. measuring cAMP levels. The EC_50_ was determined to be 39.5 ± 17.4 nM. (C) Receptor occupancy of CVN527 (p.o) measured in the striatum (n=5/group) 1 hour after dosing. (D) Terminal blood (plasma) and brain samples were simultaneously taken from the receptor occupancy animals to determine the total concentration of CVN527. The RO_50_ was calculated to be 7.2 ng/ml and 16.6 ng/mg for brain. (E) Scheme of the example recorded neuron in the DRN (top); 40x BF (bottom, left) and fluorescent (excitation 470 nm) (bottom, right) images of DRN^Vgat^ neuron with a recording pipette in acute slice. (F) Examples of membrane potential (Vm) responses of low-firing neurons to selected current injection steps (−100 pA, +60 pA, +180 pA respectively) before, during and after CVN527 bath application. (G) Dependence of AP frequency of low-firing DRN^VGAT^ neurons on injected currents from −100 pA to +180 pA and its modulation by CVN527 application (n=11; p= 0.0462 and p= 0.0379, paired t-test).

To determine the binding affinity of CVN527 against native GPR6 receptors, mouse coronal striatal sections were isolated and incubated with 2.5 nM [^3^H]-RL338 along with CVN527 over a concentration range (10^-11^ – 10^-4^ M). CVN527 displaced [H3]-RL338 with a Ki of 4.31 nM, showing potent binding to the native mouse receptor. To determine the potency of CVN527 to inhibit mouse GPR6-mediated cAMP signaling, a functional GPR6 cAMP HTRF cell-based assay was developed using CHO-K1 cells. Under these conditions, CVN527 fully inhibited GPR6 with an EC50 of 39.5 ± 17.4 nM, similar to the previously reported EC50 of 34 nM at human GPR6^28^ (**Figure 3B**). These data show that CVN527 has similar activity at human and mouse GPR6.

To correlate *ex vivo* receptor occupancy and exposure *in vivo*, wild-type mice received a single oral dose of vehicle or CVN527 (1, 3, 10, or 30 mg/kg). Plasma and striatal tissue samples were processed 1 hour after each dose (**Figure 3C**). Receptor occupancy of CVN527 at GPR6 increased dose-dependent, reaching 46% at 10mg/kg and 66% at 30 mg/kg. Plasma and brain tissue concentrations of CVN527 were quantified by LC-MS/MS bioanalysis and correlated to receptor occupancy. This revealed that the plasma or total brain concentration associated with 50% receptor occupancy was 7.2 ng/ml of CVN527 and 16.6 ng/ng, respectively (**Figure 3C**). The bioanalysis also showed a dose-dependent increase in exposure with similar brain-to-plasma ratios across doses ranging from 2.0 at 1 mg/kg to 2.6 at 30 mg/kg. These data show that CVN527 is highly brain penetrant in mice (**Figure 3D**).

We next performed slice electrophysiology to test whether CVN527 directly affects the cell excitability of DRN^VGAT^ neurons. Vgat-IRES-Cre driver mice were crossed to Cre-dependent EGFP-L10a reporter mice to selectively label vGgat-expressing cells for recordings **(Figure 3E)**. We measured changes in intrinsic AP-firing properties in response to current step injections before and after CVN527 (100nM) application. Based on their firing frequency at the baseline at any current injection step, neurons were classified as having low (<40Hz) or high (>40Hz) firing. Bath application of CVN527 reduced the firing of low-firing DRN^VGAT^ neurons, with significant values, at higher depolarization steps of 0 pA and 100 pA (n=11). **(Figure 3F-G)**. These effects were partially reversed during the washout phase. No significant effects of CVN527 were observed on high-firing neurons **(Extended Data Fig. 3)**. The reduced excitability in the low-firing group of neurons after CVN527 application is consistent with the pharmacologic finding that CVN527 is an inverse agonist for GPR6. Together, the pharmacologic and electrophysiologic studies demonstrate that CVN527 is a brain penetrant, a highly selective, and a potent inverse agonist of GPR6.

We next tested whether CVN527 treatment could replicate the effect of chemogenetic inhibition of DRN^VGAT^ neurons using two different approaches: 1) a direct infusion of 300 ng (0.1 µl/minute) of CVN527 in the DRN through an intraparenchymal brain cannula using a pump and 2) by feeding mice 10mg/kg of CVN527 mixed into a 0.2g peanut butter pellet. In both cases, the treatment was started when mice on a chow diet were switched to a HFD **(Figure 4A and 4I)**. The intra-DRN infusion of CVN527 for 14 days reduced body weight when lean mice transitioned from a chow diet to an HFD **(Figure 4B).** This effect was significantly diminished when the drug infusion was changed to vehicle (washout period).

**Figure 4.**
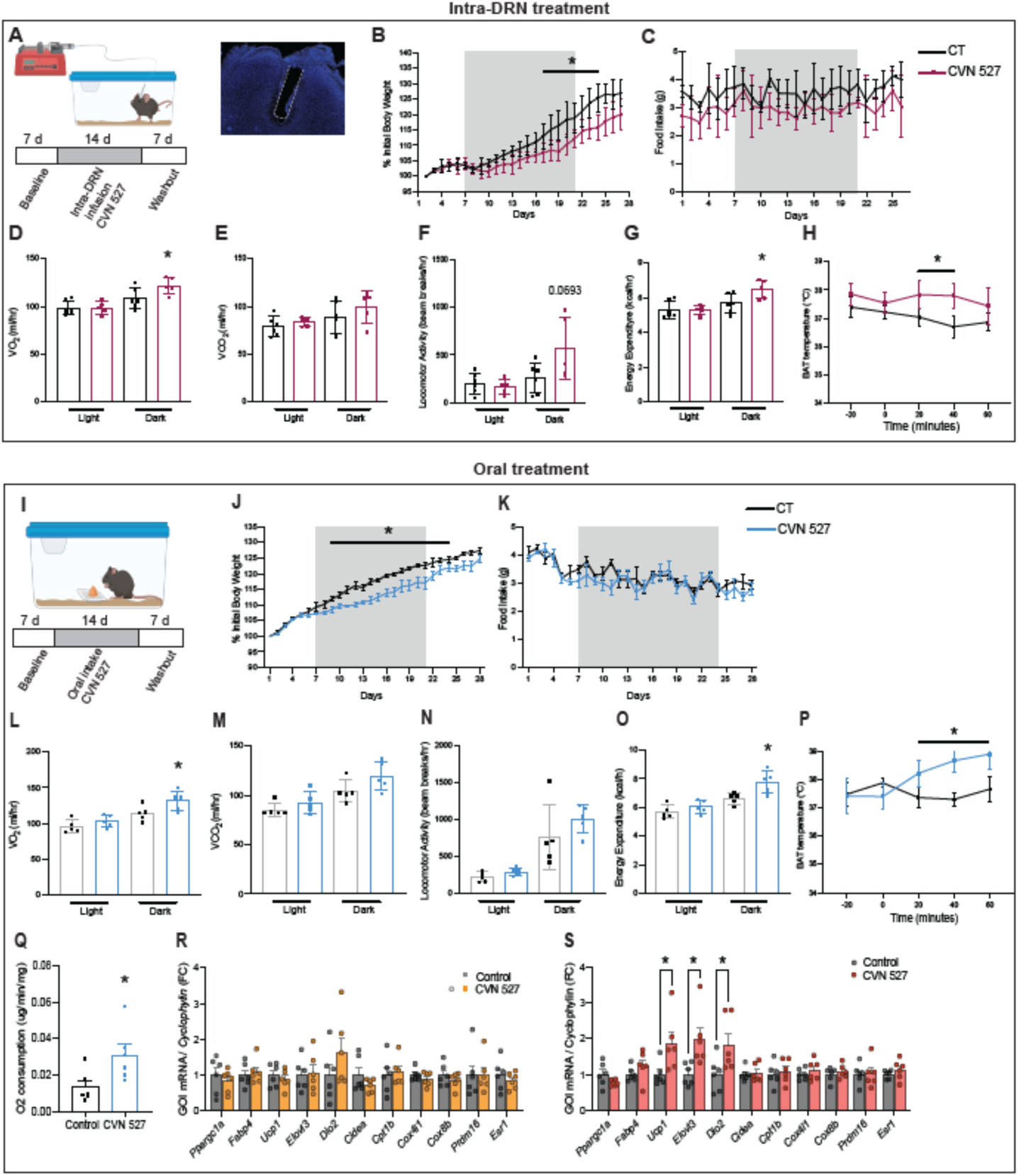
Intra-DRN and orally administrated CVN527 reduce body weight gain in DIO mice by enhancing thermogenesis. (A) Schematic for intra-DRN infusion (created with BioRender.com): following 7 days of baseline measurements and sham infusions, CVN527 was administrated for 14 consecutive days, followed by a 7-day washout period. Representative photomicrograph of cannula placement is shown on the right. (B) Weight assessment and (C) high-fat diet consumption during 28 days of the experimental period (n=5-6 per group, *P*<0.05). (D-G) Indirect calorimetry assessment using metabolic cages for evaluation of (D) oxygen consumption (VO_2_), (E) dioxide production (VCO_2_), (F) locomotion activity and (G) energy expenditure. (H) Adaptive thermogenesis after intra-DRN infusion of CVN527. (I) Schematic orally administrated CVN527: mice were given peanut butter only, following 7 days of baseline measurements, CVN527 (30mg/kg) was then mixed in peanut butter for 14 consecutive days, followed by a 7-day washout period. (K) CVN527 levels measured in plasma and brain by LC/MS/MS 30 minutes after oral administration. (L) Weight assessment and (M) high-fat-diet consumption during 28 days of the experimental period (n=5 per group, *P*<0.05). (M-P) Indirect calorimetry assessment using metabolic cages for evaluation of (M) oxygen consumption (VO_2_), (N) oxygen dioxide production (VCO_2_), (O) locomotion activity and (P) energy expenditure. (Q) Adaptive thermogenesis after oral consumption of CVN527. (R) BAT explant oxygen consumption rates detected using the Clark oxygen electrode 60 minutes after oral consumption of CVN527. (S-T) Expression of thermogenic genes (qPCR) in BAT after (S) 1 and (T) 3 days of oral consumption of CVN 527 (n=6 per group, *P*<0.05). Data are represented as mean ± s.e.m. *P* values were calculated using a two-way ANOVA with a multiple-comparisons test using a Tukey post-hoc approach (**B-C, H, J-K**), using CalR software and a two-sided ANCOVA regression analysis taking body weight into account (**D-G, L-O**) or an unpaired two-tailed Student’s *t*-test (**Q-S**). *P* < 0.05 is considered significant. The CVN527 dose used was 10 mg/kg.

Properly maintaining energy balance depends on the brain’s ability to integrate circulating hormones and fuels into major neural networks regulating feeding and thermogenesis^1,29^. Since DRN neurons have been linked to both the control of feeding behavior and thermoregulation^17,18^, we next assessed whether the reduction in the activity of DRN^VGAT^ neurons induced by intra-DRN infusion of CVN527 reduced food intake, induced thermogenesis, or both. Surprisingly, no effects on food intake were observed **(Figure 4C)**. However, when metabolic phenotyping was performed, we found increased oxygen consumption (VO2) **(Figure 4D)** and energy expenditure **(Figure 4G),** with no effect on carbon dioxide production (VCO2) **(Figure 4E)** or locomotor activity **(Figure 4F)**. To further assess if BAT might have contributed to the overall increase in metabolic rate, we implanted wireless temperature transponders (OPTT-300) into the BAT and found acute increases in the BAT temperature 20 minutes after intra-DRN infusion of CVN527 **(Figure 4H)**.

We next repeated these studies in animals receiving CVN527 orally simultaneously as animals were switched from chow to HFD **(Figure 4I).** After 14 days of treatment, we observed a significant reduction in body weight with no changes in food consumption compared with mice given peanut butter alone **(Figure 4J and K)**. Consistent with the effects we observed in the intra-DRN trial, we also observed increased oxygen consumption (VO2) **(Figure 4L)** and energy expenditure **(Figure 4O)** in mice given CVN527. No changes in carbon dioxide production (VCO2) **(Figure 4M)** and locomotor activity **(Figure 4N)** were observed. To further confirm that the effect on the prevention of weight gain was associated with the activation of BAT, we also assessed the acute effects of oral consumption of CVN527 on BAT temperature and, as with the DRN infusions, we observed a significant and persistent temperature increase after 20 minutes **(Figure 4P)**. Possibly consistent with this, patients from clinical trials with lead compound CVN424 (a different GPR6 inverse agonist) for Parkinson’s disease reported a heat sensation, and bodily temperature increases between 0.5 and 1°C were observed^30^.

The absence of GPR6 expression in BAT and the persistent thermogenic effect prompted us to further study metabolism and transcriptional changes in this tissue. Using Clark electrode, we observed increased O2 consumption in BAT 60 minutes after oral CVN527 **(Figure 4Q)**. Next, we sought to measure the expression level of canonical genes involved in the thermogenic response 1 and 3 days after CVN527 consumption. While we did not observe any significant difference in a panel of thermogenic genes 1 day after CVN527 consumption **(Figure 4R)**, we observed significantly increased expression of *Ucp1*, *Elovl3*, and *Dio2* in BAT of mice given oral CVN527 for 3 consecutive days **(Figure 4S)**. A putative explanation for the differences we found in this time-course experiment is that the BAT gene expression is highly variable depending on the pro-thermogenic stimuli^31^. For example, mice treated with CL316,243 (a selective *Adrb3* agonist) exhibit increased expression of *Ucp1* and *Pgc1a* in the inguinal adipose depot as early as one day^32^. Conversely, mice exposed to cold (6.5°C) exhibit more pronounced expression of thermogenic genes after three days^32^.

We have previously shown that reducing tonic GABAergic input from DRN onto the raphe pallidus (RPa) increases sympathetic outflow to BAT^18^. Based on this one possibility is that the increased thermogenic response we observed after CVN527 administration was driven, at least in part, by a similar mechanism involving stimulation of BAT sympathetic preganglionic neurons, resulting in increased BAT sympathetic nerve activity. Within the DRN, DRN^VGAT^ neurons locally inhibit DRN serotonergic neurons^33^. Thus, an alternative neural mechanism to explain the increase in thermogenesis response is that the disinhibition of DRN serotonergic neurons increases BAT temperature and energy expenditure upon activation^34^. Further studies will be necessary to distinguish between these and other possibilities.

Finally, to confirm the safety of CVN527, we conducted an array of behavioral tests to identify putative side effects. We first evaluated anxiety responses which often align with overt side effects. The performance of mice receiving CVN527 were evaluated in an open field test, an elevated plus maze and a marble burying experiment. Importantly, CVN527 was not associated with an increase in anxiety in any of the tests **(Extended Data Fig. 4 A-C)**. We also did not detect any aberrant behavior regarding grooming, twitching and motor behaviors using an automated system for behavioral testing **(Extended Data Fig. 4D)**.

Overall, our findings indicate that oral CVN527 dosing significantly enhances thermogenic pathways in BAT, which appears to account for the decrease in weight gain of mice transitioning to HFD. This significant reduction in weight gain underscores the potential therapeutic value of CVN527 in treating obesity.

## Acknowledgements

M.S. acknowledges support from the McCluskey family, the interstellar initiative, the ADRC Scholar fund and the Matilda Ziegler Family Foundation. M.S. and J.M.F. acknowledge support from the Robertson Therapeutic Development Fund. This work was supported by the National Institute of Diabetes and Digestive Kidney Diseases (NIDDK) grant R00DK120869 (M.S.). J.M.F. acknowledges support from the JPB Foundation. L.P. acknowledges support from The David Rockefeller fellowship program and Boehringer Ingelheim. P.C. acknowledge support from ADA Pathway to Stop Diabetes. N.R. acknowledges support from the program the ERC Consolidator grant ‘Virgins’. N.H. and J.M.F. acknowledge support from the Howard Hughes Medical Institute. We would like to thank E. Stoyanova for comprehensively organizing the GENSAT data. We would like to specially thank and dedicate this manuscript to Inna Piscitello for all her support and advice.

## Author Contributions

M.S., A.M-A.and J.M.F. conceived and designed the study and developed the research program. A.M-A, K.P., K.J.P., D.F.B., F.M., J.G.G-R., K.D., R.B., R.N., D.M., B.C., V.M.M., J.T.S., M.J.O., N.R., and L.P. performed experiments. T.S.C., D.B. and N.H. conducted GENSAT based profiling analysis. A.B., performed the electrophysiology experiments. P.C. conducted energy expenditure study design and helped select the proper panel of BAT gene expression. N.L.B., A.G. and M.C. developed the CVN compound and designed translational experiments. A.M-A., J.M.F. and M.S. wrote the manuscript with input from all the authors.

Correspondence to Alexandre Moura-Assis, Marc Schneeberger or Jeffrey M. Friedman.

## Methods

### Mice

All animal experiments were approved by the Ethical Review Committee at the local institutes and in accordance with the National guidelines on use of animals in research. For experiments conducted at Rockefeller University, mice were group housed in a 12-12h dark/light cycle at 22 °C with ad-libitum access to a regular chow diet (PicoLab Rodent Diet 205053) and water. Diet-induced obese (DIO) wild-type and Vgat-IRES-Cre male mice were fed on 45% high-fat diet (HFD, Research Diets, Cat#D12451). For all Cre mouse line experiments, only heterozygous animals were used and littermates of the same sex were randomly assigned to either experimental or control groups.

### Viral Vectors

All viral vectors used in these studies have been extensively used in neuroscience. For chemogenetic studies, AAV5-EF1a-DIO-hM4Di-mCherry (Addgene, 44632) or AAV5-EF1a-DIO-mCherry (Addgene, generated from plasmid 50462) were used. For viral-mediated translating ribosomal affinity purification (TRAP) to molecularly profile DRN^VGAT^ neurons, AAV5-IV-GFPL10 (Addgene 98704, UNC Vector Core) was used.

### Chemogenetic Studies

Mice were injected with DREADD (hM4D(Gi)) or control virus in the DRN, followed by a recovery period of at least 3 weeks (see above for viruses injected). Mice were habituated with sham injections at least 5 days prior to the assay. Mice were fed high fat diet for 16 weeks before starting the clozapine N-oxide (CNO) mediated activation of DRN^VGAT^ neurons (curative) or given high fat diet at the same time of the CNO mediated activation of DRN^VGAT^ neurons (preventive). In the curative approach food intake and body weight were measured daily after CNO injection intraperitoneally at a dose of 1mg/kg/day. In the preventive approach body weight and food intake was measured in a daily basis for 11 weeks and mice received 1 mg/kg/day of CNO twice a day (7:00 a.m. and 7p.m.). Adiposity measurements were performed using a magnetic resonance equipment (EchoMRI) system at week 1 and week 11 in the preventive setting and at day 7 and 21 in the curative setting. Thermogenesis assays were performed in the home cage uring the animal’s light phase at week 1 and week 11 in the preventive setting and at day 7 and 21 in the curative setting. Mice were given *ad libitum* access to chow diet or high fat diet during the entire experiment. BAT temperature were recorded using wireless implantable temperature probes IPTT-300 (Bio Medic Data Systems) and core body temperature was assessed using an anal probe (Braintree Scientific). Control studies were performed by injecting vehicle (saline) instead of CNO. All CNO injections were at a concentration of 1 mg/kg.

### Histology

Mice were transcardially perfused with PBS (Fisher, Cat# BP39920), followed by 4% PFA (EMS Cat # 15714-S). Brains were dissected and post-fixed in 4% PFA at 4°C for 12 hours. Brains were sectioned into 50-μm coronal slices using a Cryostat (Leica). For the immunohistochemistry reactions, brains were blocked in blocking buffer (0.2% Triton X-100 Thermo Cat# 85111 in PBS, 3% bovine serum albumin Sigma Cat# A9647-500G, 2% normal donkey serum Jackson ImmunoResearch Cat# 017-000-121) for 1-2 hours. For cFOS staining, sections were incubated with primary antibody (1:1000 rabbit anti-cFOS, Synaptic Systems 226008) overnight at 4°C. Sections were then washed 3 times in PBS and incubated with secondary antibody (1:500 donkey anti-rabbit IgG Alexa 546, Thermofisher A10040) for 2 hours at room temperature, washed 3 times in PBS and mounted with DAPI Fluoromount-G (Southern Biotech Cat# 0100-20). Images were taken using a LSM780 confocal microscope (Zeiss) and ImageJ software (NIH) was used for minimal processing including brightness and contrast adjustments. cFOS counts were conducted manually using image j in 4-5 DRN sections of 5 animals from anterior posterior regions spanning −4.1 to −4.8 mm.

### Multiplex fluorescence *in-situ* hybridization (FISH) and hybridization chain reaction (HCR)

Mice were transcardially perfused with RNAse-free PBS, followed by sterile 4% PFA. Harvested brains were subsequently frozen and sectioned into 20-50μm coronal slices using a Cryostat (Leica). Multiplex FISH was then performed using *Slc32a1* and *gpr6* probes (Advanced Cell Diagnostics) for targeting its correspondent mRNAs. Formalin-fixed frozen brain tissue was sectioned using a cryostat. The sections obtained were attached on Superfrost Plus Adhesion Slides (Thermo Fisher), and a hydrophobic barrier created using Immedge Hydrophobic Barrier Pen (Vector Laboratories Inc.). Pre-treatment was done by serial submersion of the slides in 1X PBS, nuclease-free water, and 100% EtOH for two minutes each, at RT. Probe hybridization was achieved by incubation of 35 µL mRNA target probes for 2 h at 40°C using a HyBez oven. The signal was amplified by subsequent incubation of Amp-1, Amp-2, and Amp-3; one drop each for 30, 30, and 15 min respectively at 40°C using a HyBez oven. Each incubation step was followed by two 2 min washes using RNAscope washing buffer. Nucleic acids were stained using manufacturer’s supplied DAPI for 30 s followed by two washes with 1X PBS. The slides were cover slipped and mounted using Prolong Gold Antifade Mountant (Thermo Fisher).

### Human tissue analysis

The expression of GPR6 mRNA in raphe nuclei from human subjects was evaluated in frozen sections of midbrain from two non-diseased male donors. Fully consented, post-mortem human tissues were obtained from Tissues for Research (UK). Duplex RNAscope assays (Advanced Cell Diagnostics, Biotechne) were completed using probes to SLC32A1 (c1) and GPR6 (c2). On completion of the assays, sections were scanned in bright-field at 40x (Hamamatsu Nanozoomer) and analyzed for the presence and distribution of single or double-labeled raphe neurons.

### Stereotaxic surgery

Mice were anesthetized with isoflurane (3% induction, 1.5-2% maintenance) in oxygen and placed in a stereotax apparatus (Kopf Instruments). Viruses were delivered into the DRN through a glass capillary using a Drummond Scientific Nanoject III. For the DRN coordinates, the following coordinates relative to the lambda: −2.8mm DV, 0 mm ML, 0 mm AP). For pharmacology studies, wild-type mice had a cannula (Plastics One) placed in the DRN (coordinates, relative to lambda: +0.8 mm ML, 0 mm AP, −3.0 mm DV:15°).

### Intra-DRN administration of compound CVN527

Drug infusions were achieved employing an infusion pump (Harvard Apparatus) attached to glass syringes (Hamilton), with an infusion rate of 0.1 µl/minute. Total drug injection never exceeded 0.5 µl in local DRN studies for a total of 300ng. Mice were habituated for 1 week to be plugged to the infusion system and minimize stress. After habituation mice were measured (body weight and food intake) during 7 days in a vehicle phase. Next, mice were divided in aged-matched and weight-matched groups. The first group of mice was treated with CVN527 and the second group of mice received only vehicle without the CVN527 compound. After the 14 treatment days, mice returned for 7 days on vehicle only to assess whether there were irreversible consequences in weight due to the chronic treatment.

### Oral administration of compound CVN527

Cerevance Therapeutics proprietary preclinical specific oral GPR6 inverse agonist CVN527 has been administered at a dose of 10mg/kg daily mixed in a pellet of 0.1g of peanut butter in mice fed for 16 weeks in HFD until they reach DIO. Mice were habituated 1 week prior to the study to receive 1 pellet of peanut butter. All mice successfully ate the pellet for first day of habituation in less than 2 minutes. After habituation mice were measured (body weight and food intake) during 7 days in a vehicle phase while only receiving peanut butter. Next, mice were divided in aged-matched and weight-matched groups. The first group of mice was treated with CVN527 and the second group of mice received only the peanut butter pellet without the CVN527 compound. Treatment lasted 14 days to mimic DRN treatment. A lean control group was also evaluated in parallel following the same paradigm. After the 14 treatment days, mice returned for 7 days on peanut butter only to assess whether there were irreversible consequences in weight due to the chronic treatment.

### iDISCO+ and ClearMap analysis

Whole-brain staining was performed following the iDISCO+ protocol previously described^16^. Briefly, perfused brains were dehydrated in methanol 20%, 40%, 60%, 80% and 100% (Sigma-Aldrich) and incubated overnight in a 66% dichloromethane solution (Sigma-Aldrich) in methanol. Samples were bleached overnight at 4°C in methanol containing 5% hydrogen peroxide (Sigma-Aldrich) and then rehydrated after incubation in methanol 60%, 40% and 20%. After permeabilization, samples were incubated with rabbit cFOS antibody, 1:4000 (Synaptic Systems 226 008) at 37°C for 10 days. Secondary antibodies conjugated to Alexa 546 were used (Life Technologies). After a final incubation in methanol for dehydration, brain samples were washed in dichloromethane 100% for clearing and stored in dibenzyl ether (Sigma-Aldrich). The imaging acquisition was performed in a light sheet microscope (LaVision Ultramicroscope II) equipped with sCMOS camera and LVMI-Fluor x4 objective lens equipped with a 6 mm working distance dipping cap. Images were taken every 6 μm (z steps) and numerical aperture set to 0.03NA. Reconstructed images were further processed with ClearMap2 software (https://github.com/ChristophKirst/).

### TRAP

Mice were sacrificed by cervical dislocation and a ventral piece including the DRN was rapidly dissected in ice-cold buffer containing 10 mM HEPES [pH 7.4], 150 mM KCl, 5 mM MgCl2) with 0.5 mM DTT (Sigma), 80 U/ml RNasin Plus (Promega), 40U/ml SuperaseDIn (Life Technologies), 100 mg/ml cycloheximide, protease inhibitor cocktail (Roche) and then cleared by centrifugation to isolate polysome-containing cytoplasmic supernatant. Polysomes were immunoprecipitated using monoclonal anti-EGFP antibodies (clones 19C8 and 19F7 bound to biotinylated-protein L-coated streptavidin-conjugated magnetic beads (Pierce, Thermo Fischer; Life Techonologies). A small amount of RNA was taken prior the immunoprecipitation (Input) and both the Input and immunoprecipitated (IP) were purified using the Absolutely RNA Nanoprep Kit (Agilent). For RNA-seq, cDNA libraries were prepared with the SMARTer Ultralow Input RNA for Illumina Sequencing Kit (Clontech) and sequenced on an Illumina HiSeq2500 platform.

### RNA-seq analysis

Sequence and transcript coordinates for mouse mm10 UCSC genome and gene models were retrieved from the Bioconductor Bsgenome. Mmusculus.UCSC.mm10(version1.4.0) and TxDb. Mmusculus. UCSC. mm10. Known Gene (version 3.4.0) Bioconductor libraries, respectively. FASTQ files for the published DRNVgat neuron TRAP experiments were obtained from the Gene Expression Omnibus (accession: <u>GSE87890</u>). Transcript expressions were calculated using the Salmon quantification software^35^ (version 0.8.2) and gene expression levels as TPMs and counts were retrieved using Tximport (version 1.8.0). Normalization and rlog transformation of raw read counts in genes were performed using DESeq2 ^36^ (version 1.20.0). For visualization in genome browsers, RNA-seq reads are aligned to the genome using Rsubread’s subjunct method (version 1.30.6) and exported as bigWigs normalized to reads per million using the rtracklayer package^37^ (version 1.40.6). GSVA analysis of selected genes was performed using the GSVA (version 1.34.0) R package. Visualization of genes and gene-sets as heat maps was performed using the Pheatmap R package (version 1.0.10).

### GENSAT-TRAP comparison

Gene expression enrichments of the immunoprecipitants from Vgat3 neurons were compared with expression data in the GENSAT database previously processed as described above (see ‘RNA-seq analysis’). Adjusted *P* values for the Vgat neurons (TRAP IP samples) versus each individual experiment in the GENSAT database were calculated with DESeq2. A combined *P* value and chi-squared statistic for all of these comparisons for each gene were then calculated using Fisher’s method, specifically with the metap R package (version 1.4). The gene list was filtered further to those that are in both the signaling receptor activity (<u>GO:0038023</u>) and plasma membrane (<u>GO:0005886</u>) Gene Ontology categories. Selected receptors for further study were validated using ISH, and the expression of these receptors compared to the GENSAT samples was also visualized using the Complex Heatmap Bioconductor R package^38^.

### Home-Cage indirect calorimetry assessment

Energy expenditure was assessed through measurement by indirect calorimetry of oxygen consumption (VO2), carbon dioxide production (VCO2), heat production, and locomotor activity using an automated home cage phenotyping system (TSE Systems). Mice were singled housed in a climate-controlled chamber (temperature: 22°C; humidity: 55%; 12-hour light/12-hour dark cycle) with ad libitum access to water and chow. After 7 days (TSE Systems) of adaptation to social isolation, the mice were treated with vehicle (4 days) or CNO (6 days) **(Figure 1)**, or peanut butter (0.1g) with (6 days) or without (4 days) CVN527. Data were collected and analyzed as recommended by the manufacturers. Averaged measurements of control periods or treatment periods (CNO/CVN527) are represented. Locomotor activity was recorded as beam breaks converted into distance/velocity, measuring activity in 3D. Beam breaks by mice were analyzed in the metabolic cage using custom software. The respiratory exchange ratio (RER) and energy expenditure were calculated from VO2 and VCO2 production data. Statistical assessment of the data was conducted using CalR software^43^ and Ancova regressions.

### RNA extraction and BAT gene expression analysis

Intrascapular BAT was extracted 1-hour post-treatment; white adipose tissue was carefully trimmed and BAT snap frozen. Brown fat pads were pulverized in liquid nitrogen and tissue powder was split for RNA and soluble protein extractions. RNA was obtained using a combined TRizol (Life Technologies, Cat. 15596018) and RNeasy spin column (Qiagen, Cat.74104) protocol. A total of 2ug RNA was used for cDNA synthesis (Thermo Scientific, Cat. 4368813) per sample. Expression of *Ucp1*, *Elovl3*, *Cidea*, *Cox8b*, *Ppargc1a*, *Esr1*, *Dio2*, *Cpt1b*, *Cox4l1*, *Fabp4* and *Prdm16-ex9* (exon 9) was assessed with Syber Green (Applied Biosystems, Cat. 4367659) in 384-well plate. Cyclophylin was used a reference gene for delta analysis. Forward and reverse primers sequences are listed below:

#### qPCR primers

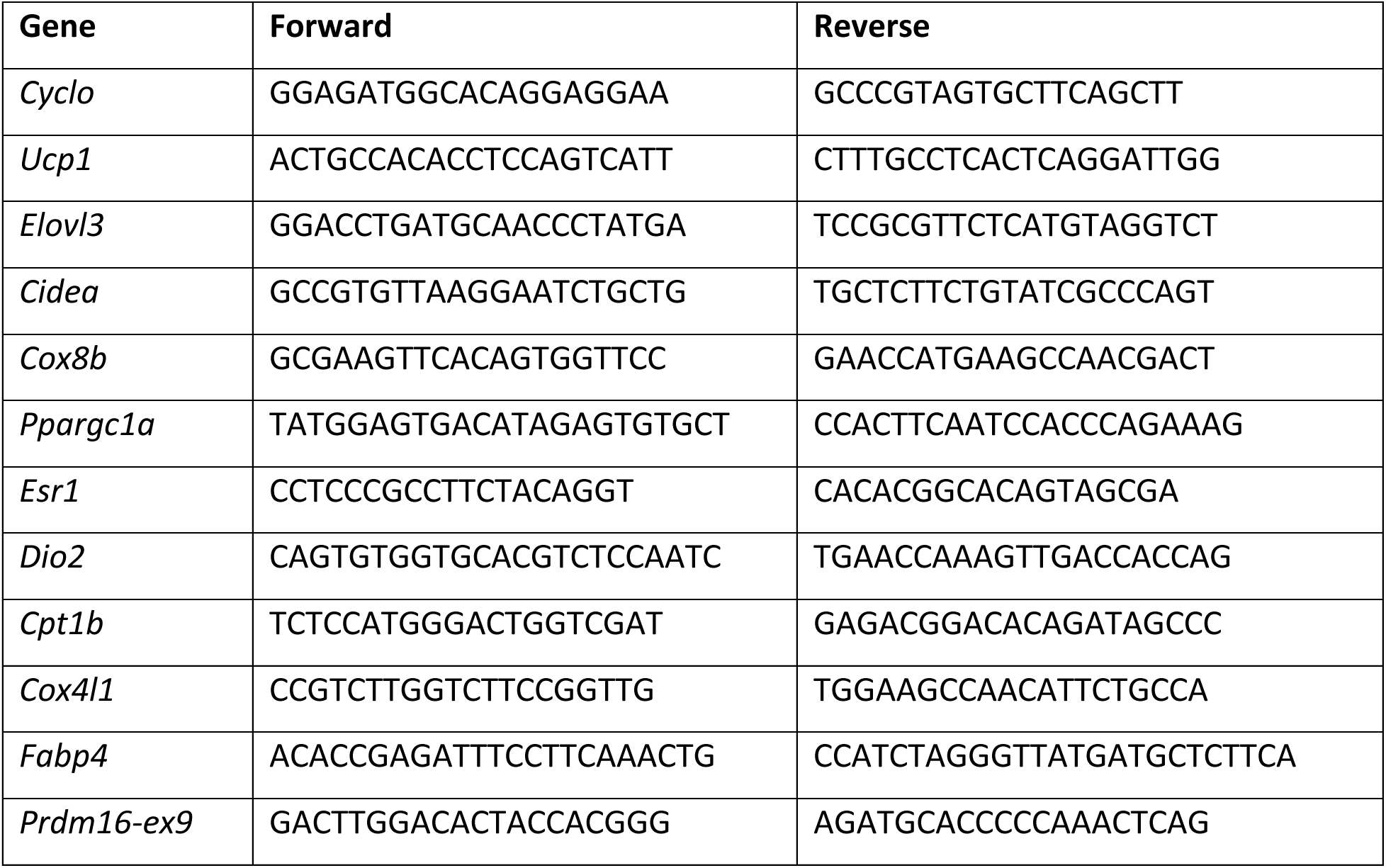

### Brown adipose respiration in Clark Electrode

Oxygen consumption was assessed using Clark Electrode (Si Strathkelvin Instruments, Mitocell MT200) in freshly dissected intrascapular BAT from mice 1hr-post GPR6 treatment. Polypropylene electrodes were first washed with acetone then electrode jackets were filled with KCl and bubbles were removed to ensure electrode’s membrane was watertight. Electrodes were connected to respirometry chamber and left in H_2_O overnight to stabilize to 0 pA. Oxygen solubility at 37°C for salinity of 0 and a pressure of 760 mmHg was set to 6.73 mg/L. Electrodes were then calibrated using pre-warmed (37°C) O_2_-low (0.25% Na_2_SO_3;_ pA 0-8) and O_2_-high (oxygen-saturated H_2_O; pA 600). One sample was tested at a time in 15-min intervals. BAT was dissected, placed in saline on ice and immediately divided into 3 pieces (50-100mg each). Each tissue piece was weighed using analytic scale, minced with spring scissors (40 snips) and buffered in 0.5mL of respiration buffer (DPBS, 20% BSA, 4.5% glucose, 1mM Na pyruvate). Resuspended tissue pieces were immediately placed in the respiration chamber with a stirring bar, and O_2_ consumption rates were recorded by the electrode for 2 mins each. Respiration chamber was rinsed with water and fresh respiration buffer between samples. Consumption rates were normalized to tissue weights and auto calculated by the Si Strathkelvin software. iBAT temperature was measured using wireless implantable temperature probes IPTT-300 (Bio Medic Data Systems), and core body temperature was assessed using an anal probe (Braintree Scientific).

### Patch-clamp electrophysiology

Animals were anesthetized with 4% isoflurane for 5 minutes until reflexes are not present and then decapitated (according to the authorized protocol). 250 µm thick coronal brain slices containing the dorsal raphe nucleus (DRN) were made with a Leica VT1000S vibratome (Leica Microsystems, Wetzlar, Germany). DRN was identified by the presence of aqueduct. Slicing was done in a high sucrose solution containing (in mM): 87 NaCl, 2.5 KCl, 1.25 NaH2PO4, 26 NaHCO3, 10 D-glucose, 50 Sucrose, 3 MgCl2, 0.5 CaCl2 (pH 7.3). Slices were maintained in a holding chamber containing extracellular solution for 45 min at 35C. Whole-cell patch clamp recordings were performed using an extracellular solution containing (in mM): 125 NaCl, 2.5 KCl, 1.25 NaH2PO4, 25 NaHCO3, 25 D-glucose, 0.1 MgCl2, 3 CaCl2 (pH 7.4, continuously bubbled with 95% O2 / 5% CO2). Recording pipettes were made from borosilicate glass (Sutter Instrument) and had resistances of 5–6 MOhm when filled with a pipette solution containing (in mM): 135 Potassium Methyl Sulfate (KCH3SO4), 4 Mg-ATP, 0.3 Na-GTP, 8 NaCl, 10 HEPES, 5 Na2-P-creatine; pH 7.31, adjusted with KOH. Whole-cell current-clamp recordings were obtained under visual control using a 40x objective using EPC-10 patch clamp amplifier under the control of PatchMaster software (HEKA Elektronik, Reutlingen, Germany). The recordings were done at room temperature. Brain slices were imaged using a CMOS camera (C11440-36U; Hamamatsu Photonics, Hamamatsu City, Japan) under control of HOKAWO 3.0 software. To visualize the eYFP fluorescence, the fluorophores were excited using LEDs (465 nm, Green; CoolLED pT-100). One current injection step round consisted of 1s steps ranging from −100 to +180 pA in 20 pA increments relative to the holding current. CVN527 was dissolved in DMSO with final bath perfusion concentration of DMSO 0.01% and CVN527 100 nM. Baseline of 5 min of current injection steps was recorded. Subsequently, 0.01% DMSO alone followed by a drug or direct drug application was performed. After 10-20 min of the drug application in the slice the washout was started. Electrophysiological recordings were analysed in IgorPro 7 (WaveMetrics) software using the NeuroMatic plug-in^39^. Cells with a leak current more than 300 pA were rejected. APs were detected using automated selection where the threshold of −10mV was manually selected. In the **Figure 3G** firing rates of 2 current injection step rounds were averaged for baseline; 2 current injection step rounds 3 min after the drug application and 2 current injection step rounds 3 min into washout were averaged.

### In vitro Pharmacology

Functional activity assessment of inverse agonists against GPR6 has been previously described ^40^ Briefly, CHO-K1 cells (ATCC cat #CCL-61) stably transfected with mouse GPR6 cDNA (Origene) were incubated in 96 well plates for 24 hours at 37◦C before being assessed via functional LANCE HTRF cAMP assay system (Perkin Elmer). CVN527 was serially diluted in DMSO and added to the plates in Kreb’s Ringer’s buffer at 1x concentration (20 μl/well) and incubated or 45 minutes at 37◦C. Final concentrations ranged from 30 µM to 0.03 nM. In a final step, Eu-labeled cAMP tracer (10 µl) dilated in cAMP detection/lysis buffer with 10 µl ULight-anti-cAMP diluted in the same buffer was added to the cells. After a 60-minute incubation at room temperature, time resolved energy transfer was measured on an EnVision plate reader (Perkin Elmer). CVN527 showed full inverse agonism (Emax 100%) and efficacy values were calculated by fitting data to the four-parameter variable slop model without constrains using non-linear regression in Prism (Graphpad).

### Receptor occupancy, Ki determination and Pharmacokinetics

*Ex vivo*, dose-dependent receptor occupancy was determined in male C57/Bl6 mice (Charles River, UK). Since receptor occupancy is expected to be related to exposure and this ratio not to be sex dependent, we chose to use one sex in these studies, furthermore this relates to the HFD and behavior studies.

The mice were dosed with vehicle (10% DMSO, 0.5% tween80, 89.5% PBS) or CVN527 at 1, 3, 10, or 30 mg/kg p.o (n=5/group) and then euthanized after 1 hour. Terminal blood samples were taken by cardiac puncture and placed in to lithium/heparin tubes (Sarsedt). Blood samples were centrifuged (1900g for 5 minutes) to extract the plasma which was transferred to a microtube and frozen at −80◦C until bioanalysis. The brains were excised, and a coronal section was taken for receptor occupancy (rapidly frozen in isopentane cooled to−20◦C), and the remaining brain tissue was stored at −80◦C until bioanalysis.

The brain section containing the striatum was cut into 20 µm sections and placed in humified chambers for 120 minutes in 200 µl buffer (50mM Tris buffer, pH 7.4 containing 50mM NACl, 6 mM MgCl_2,_ and 0.1% BSA) containing either 2.5nM [^3^H]-RL338 (total binding) and CVN527 at four different doses (1, 3, 10, 30 mg/kg) or 2.5nM [^3^H]-RL338 and 10 µM CVN527 (non-specific binding). After incubation, the slides were washed 3x in ice-cold assay buffer and 1x distilled water. Slides were analyzed on a β-imager.

Occupancy (*O*) data analysis was determined by the equation below^41^. In brief, the percentage *O* of test compound-treated animal was calculated according to the following equation:

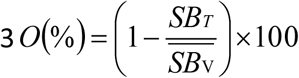

in which *SB_T_* is the specific binding in treated slices and *SBv* is the mean specific binding of the vehicle group.

Brain samples for bioanalysis were homogenized in 0.1M PBS before further processing. Both brain and plasma samples were analyzed via LC-MS/MS.

The Ki of CVN527 for mouse striatal expressing GPR6 was determined using a similar method to the receptor occupancy, except the mice were not dosed with CVN527 prior to euthanasia, and the sections (8/animal, n=9 mice) were incubated with one of nine concentrations of CVN527 (10-11 to 10-4 M). The binding (CPM/mm2) was used to determine the Ki using the Cheng-Prusoff equation^42^.

### Behavioral studies

The OFT was used to evaluate locomotion activity after treatment with CVN527 (10mg/kg) in peanut butter or peanut butter alone as control. Standard OFT arenas were used. Of note, mice had never been exposed to the OFT arena prior to studies but had been thoroughly habituated to the room where behavioral experiments occurred. Experiments were performed in 5-minute sessions (pre and post drug). Total distance traveled, velocity and time spent in the center versus time spent in the borders of the OFT arena (anxiety-related behaviors) were calculated by using an automated tracking system (Ethovision XT).

The EPM was used to evaluate anxiety after treatment with CVN527 (10mg/kg) in the same experimental groups aforementioned. Standard EPM arenas were used. Of note, mice had never been exposed to the EPM arena prior to studies but had been thoroughly habituated to the room where behavioral experiments occurred. Experiments were performed in 5-minute sessions. Total distance traveled, frequency and cumulative duration in the open arms versus closed arms, velocity and mean distance to each arm were evaluated.

The marble burying test was used to assess anxiety, twelve glass marbles were evenly spaced in 3 rows, on an approximately 5 cm layer of sawdust bedding lightly pressed down to make a flat, even surface, in a plastic cage. Each mouse was placed in the cage and left for 30 minutes, after which the number of marbles buried with sawdust was counted as a measure of anxiety.

For a battery of observational side effects (moving right, moving left, grooming, twitching, sleep eating and drinking), mice were removed from their home cage and placed individually in clear glass observation cages (36 cm × 20 cm × 20 cm). Behavioral assessments were carried out in a manner similar to that described previously for dopamine receptor D1A, D2, and D3 mutants using a rapid time-sampling behavioral checklist. The assessment cycles occurred over a 20-hour period (0–20 hours). Under these conditions, each animal was observed on one occasion only, with all assessments made by an observer who was unaware of the genotype of each animal.

### Quantification and statistical analysis

Statistical parameters reported in the Figures and Figure Legends are displayed as mean ± SEM. Significance was defined as p < 0.05. Significance annotations are annotated with the actual value. Mice were randomized into control or treatment groups. Control mice were age-matched littermate controls where possible. All statistics and data analysis were performed using GraphPad Prism, CalR Matlab, R, or Python. For RNA-seq, transcript abundance and differential expression were performed using cufflinks.

---

**Extended Data Fig.1.**
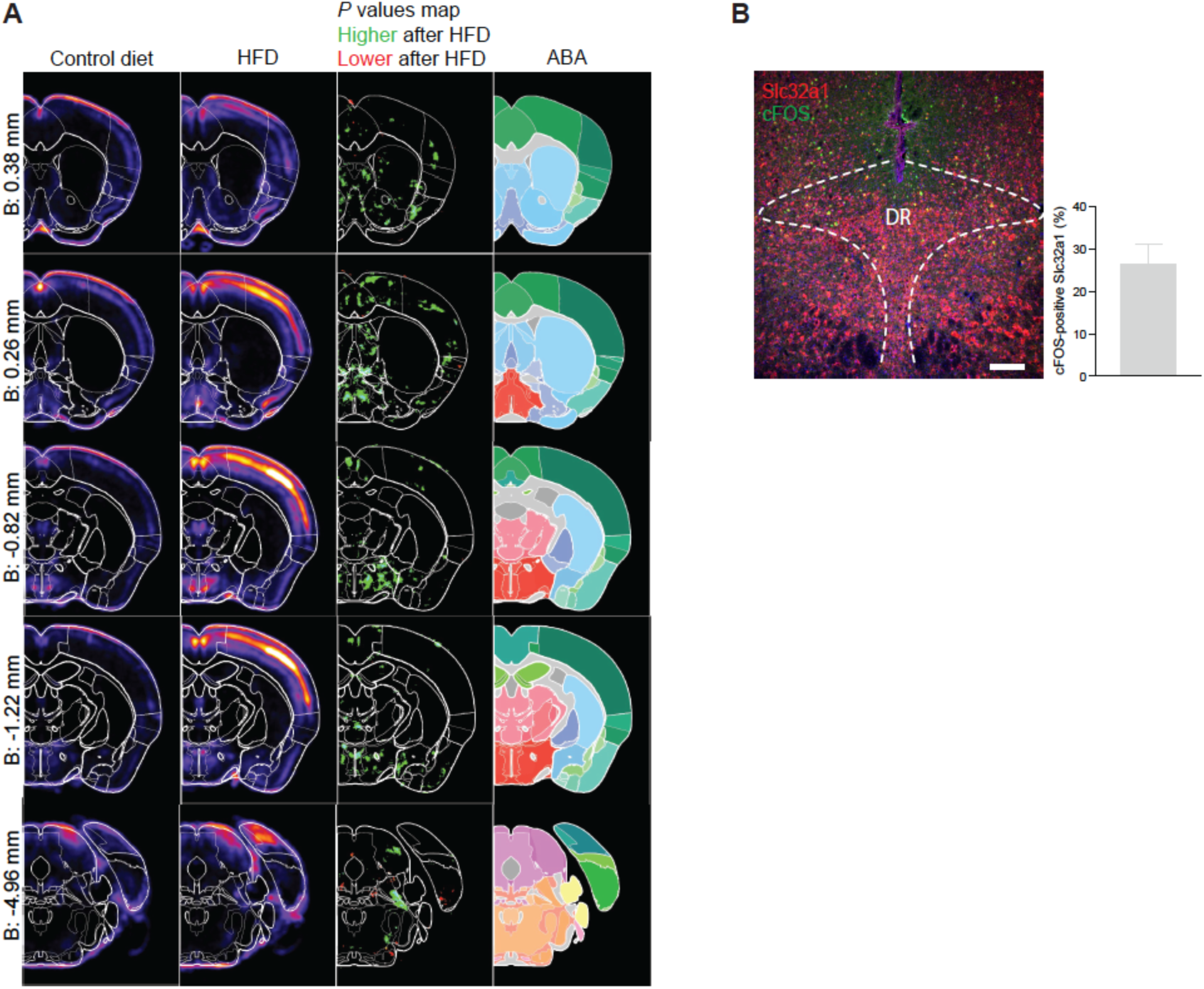
(A) Heatmaps and coronal views of the p value of the voxel maps for Fos-positive cell densities at different levels of mice brain. Regions significantly activated after 7 days of HFD feeding are highlighted in green. ABA (Allen Brain Atlas) annotated image. (B) Representative photomicrographs of immunohistochemical staining of cFOS and *Slc32a1* fluorescent *in situ* hybridization in the DRN of mice fed a HFD for 16 weeks. DR, dorsal raphe. (n=2 samples, scale bar = 100um)

**Extended Data Fig.2.**
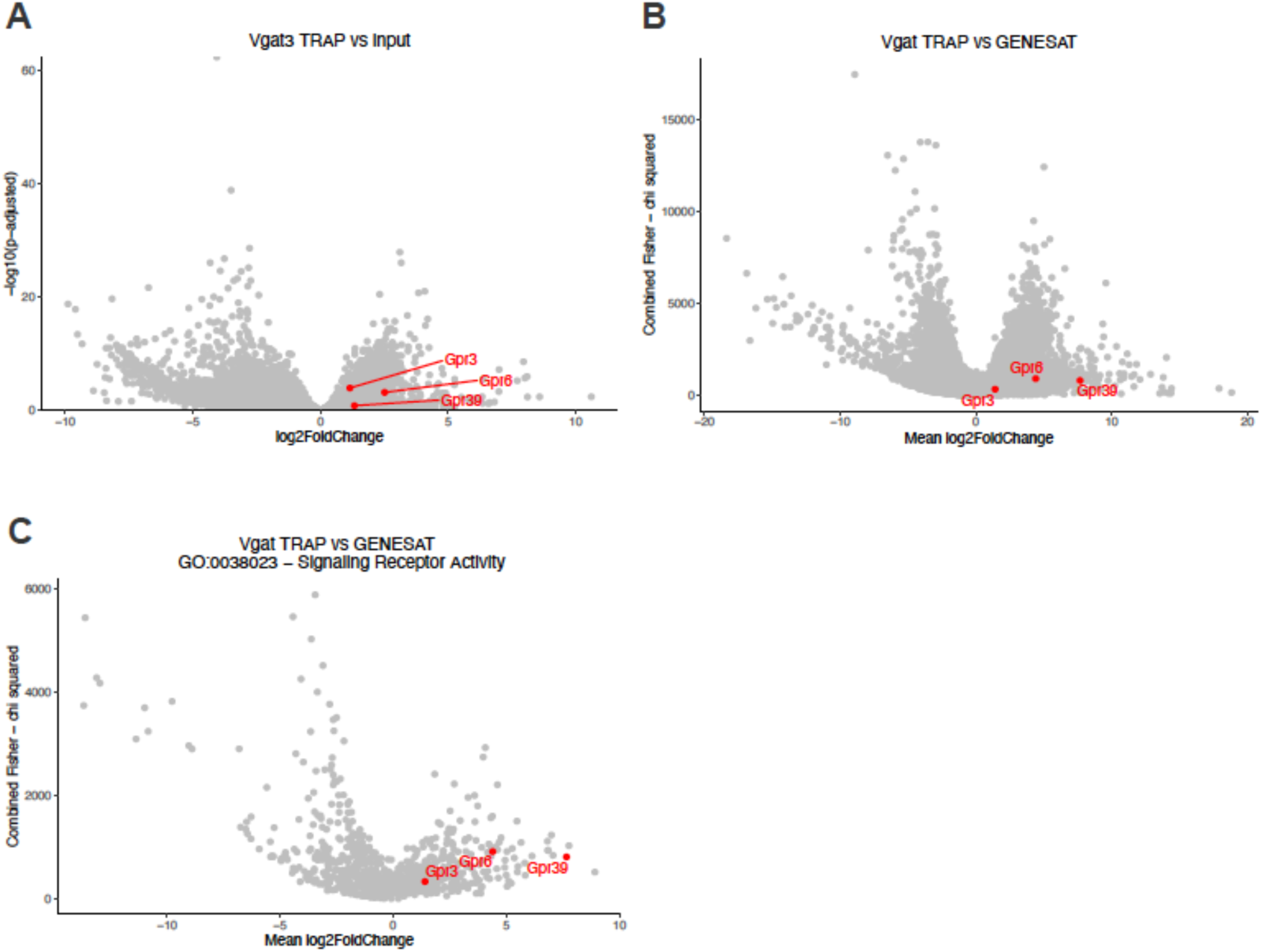
(A) Volcano plot for the adjusted p-value logarithmic plots versus the fold change logarithmic plots illustrating the set of markers most enriched in the IP sample versus the input. Red dots highlight differentially enriched markers for drug discovery that will be used to illustrate the enrichment after each filter is applied (GPR6, GPR3 and GPR39). (B) Volcano plot for the adjusted p-value logarithmic plots versus the fold change logarithmic plots illustrating the set of markers most enriched in the IP sample versus the GENSAT database. Red dots highlight differentially enriched markers for drug discovery that will be used to illustrate the enrichment after each filter is applied (GPR6, GPR3 and GPR39). (C) Volcano plot for the adjusted p-value logarithmic plots versus the fold change logarithmic plots illustrating the set of markers most enriched in the IP sample versus the GENSAT database and after applying the Gene Ontology filter of plasma membrane receptors. Red dots highlight differentially enriched markers for drug discovery that will be used to illustrate the enrichment after each filter is applied (GPR6, GPR3 and GPR39). Data are represented as enrichment in IP over input.

**Extended Data Fig.3.**
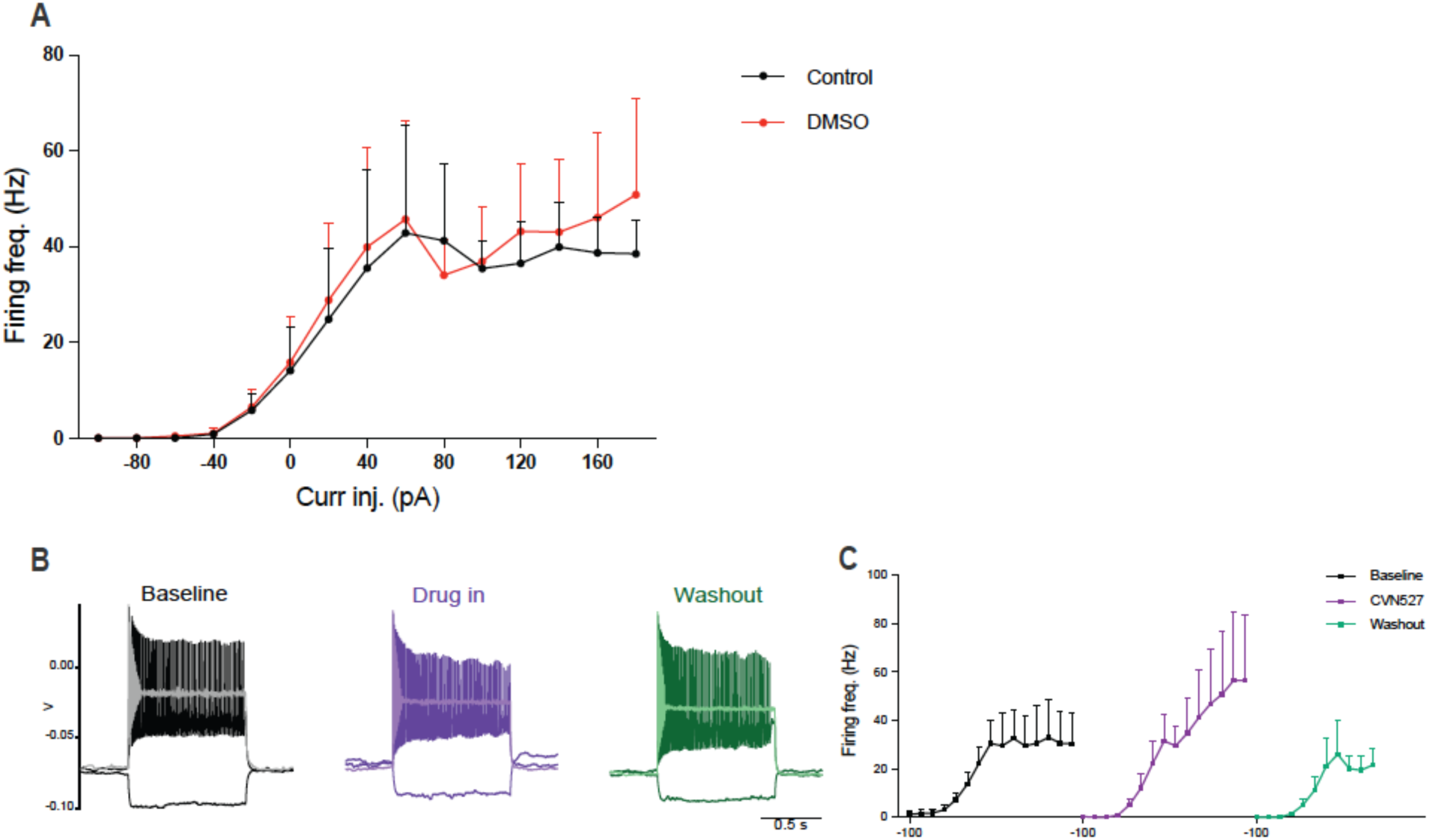
(A) Dependence of AP frequency of DRN^VGAT^ neurons on injected currents from −100 pA to +180 pA before and after 0.01% DMSO bath perfusion (n=5). (B) Examples of membrane potential (Vm) responses of high-firing neurons to selected current injection steps (−100 pA, +60 pA, +180 pA respectively) before, during and after CVN527 bath application. (C) Dependence of AP frequency of high-firing DRN^VGAT^ neurons on injected currents from −100 pA to +180 pA and its modulation by CVN527 application (n=4).

**Extended Data Fig.4.**
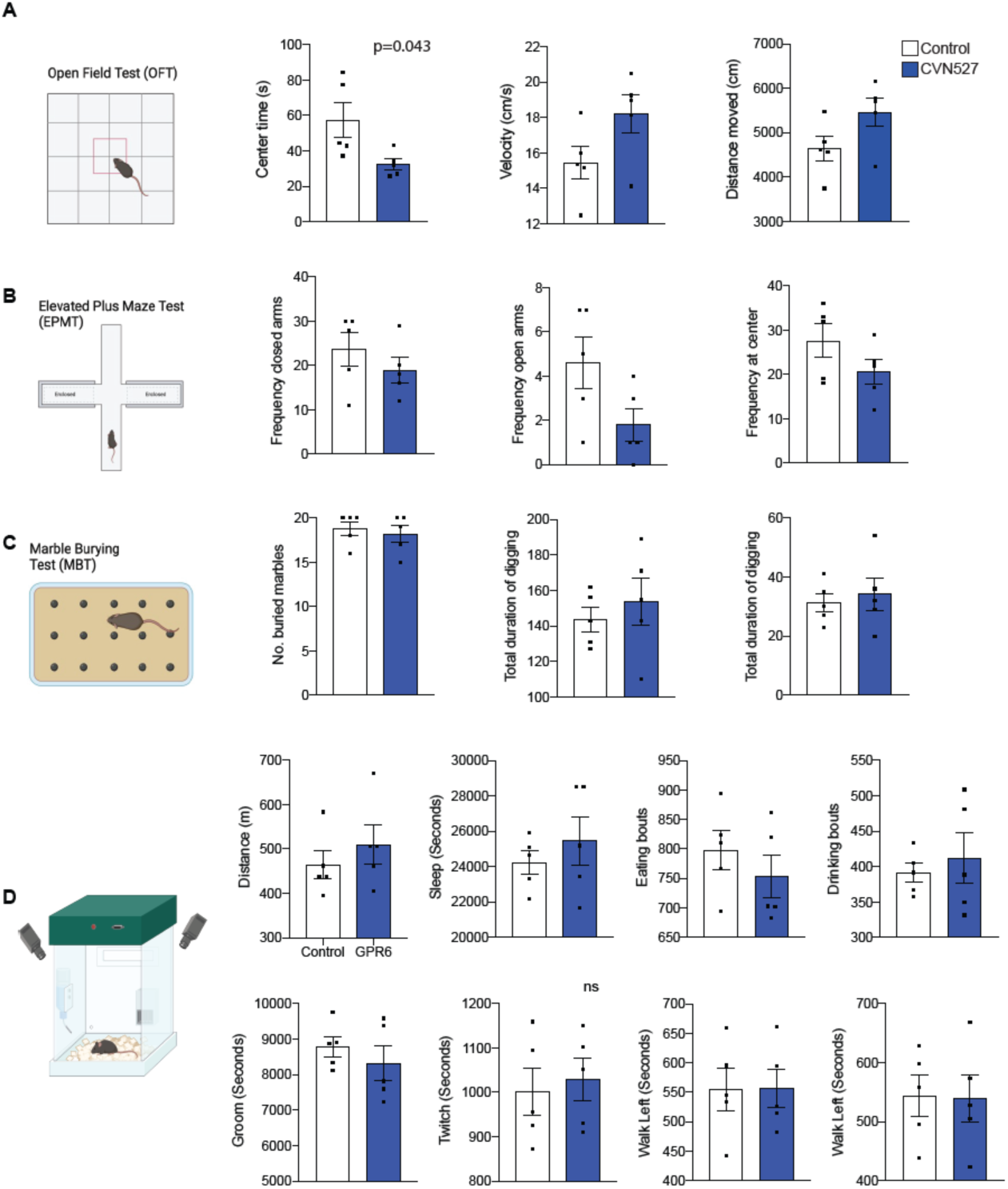
(A) An Open Field Test (OFT) is used to assess locomotion activity and anxiety. A reduction of time spent in the center (measure of anxiety) of the OFT is observed in CVN527 (10mg/kg) treated mice arguing for a lower level of anxiety. A locomotor effect is not observed in terms of total distance travelled and velocity in a 5 min session. (B) An elevated plus maze (EMP) is used as an established protocol for anxiety. No differences are observed in the time spent in the closed arms (left), open arms (middle) or center (right) of the EPM further suggesting no effect in anxiety at a dose of CVN527 (10mg/kg). (C) A marble burying test, in which fifteen glass marbles are evenly spaced in 3 rows shows no differences in number of marbles buried or time spent digging/non-digging suggesting no major stress or depressive effect of CVN527 (10mg/kg). (D) 20-hour automated cage observation of mice behavior in a clear glass cage to evaluate whether CVN527 (10mg/kg) lead to debilitating side effects (n = 4 per group). Behavioral paradigms evaluated are time spent ingesting food, drinking and sleeping. In addition, automated system evaluates the locomotion activity of mice as time spent walking left, walking right. As a function of that parameter, it provides the Total Distance Travelled. Recordings provide information regarding time spent grooming or twitching in the behavioral cage which would be associated to stress behaviors. Data are represented as mean ± s.e.m. *P* values were calculated using an unpaired two-tailed Student’s *t*-test. P<0.05 is considered significant.

